# Two ligand-binding sites on SARS-CoV-2 non-structural protein 1 revealed by fragment-based x-ray screening

**DOI:** 10.1101/2022.06.12.495816

**Authors:** Shumeng Ma, Shymaa Damfo, Jiaqi Lou, Nikos Pinotsis, Matthew W. Bowler, Shozeb Haider, Frank Kozielski

**Author notes:** **Corresponding author:** Frank Kozielski, Department of Pharmaceutical and Biological Chemistry, School of Pharmacy, University College London, 29-39 Brunswick Square, London WC1N 1AX, UK.

## Abstract

The regular reappearance of coronavirus (CoV) outbreaks over the past 20 years has caused significant health consequences and financial burdens worldwide. The most recent and still ongoing novel CoV pandemic, caused by Severe Acute Respiratory Syndrome coronavirus 2 (SARS-CoV-2) has brought a range of devastating consequences. Due to the exceptionally fast development of vaccines, the mortality rate of the virus has been curbed to a significant extent. However, the limitations of vaccination efficiency and applicability, coupled with the still high infection rate, emphasise the urgent need for discovering safe and effective antivirals against SARS-CoV-2 through suppressing its replication and or attenuating its virulence. Non-structural protein 1 (nsp1), a unique viral and conserved leader protein, is a crucial virulence factor for causing host mRNA degradation, suppressing interferon (IFN) expression and host antiviral signalling pathways. In view of the essential role of nsp1 in the CoV life cycle, it is regarded as an exploitable target for antiviral drug discovery. Here, we report a variety of fragment hits against SARS-CoV-2 nsp1 identified by fragment-based screening via X-ray crystallography. We also determined the structure of nsp1 at atomic resolution (0.95 Å). Binding affinities of hits against nsp1 were determined by orthogonal biophysical assays such as microscale thermophoresis and thermal sift assays. We identified two ligand-binding sites on nsp1, one deep and one shallow pocket, which are not conserved between the three medially relevant SARS, SARS-CoV-2 and MERS coronaviruses. Our study provides an excellent starting point for the development of more potent nsp1-targeting inhibitors and functional studies on SARS-CoV-2 nsp1.

## Introduction

In the past two decades, coronaviruses (CoVs) have posed a constant threat to humans in various forms, from Severe Acute Respiratory Syndrome Coronavirus (SARS-CoV), the etiological agent for the outbreak of severe acute respiratory syndrome (SARS) in 2002 [1], to the Middle East Respiratory Syndrome Coronavirus (MERS-CoV) in 2012 [2], and the most recent novel Severe Acute Respiratory Syndrome Coronavirus 2 (SARS-CoV-2), leading to the ongoing coronavirus pandemic first emerging at the end of 2019 in Wuhan, China [3]. The high infectivity and fatality of CoVs have caused great public concern [4] and its worldwide impact is causing a huge financial burden [5-7]. These events stress the importance of timely precaution and effectively curative strategies such as vaccination and the development of antiviral agents, which significantly curbed the effects of the disease [8, 9]. However, the vaccination efficacy partially depends on the physical [10] and psychological [11] conditions of inoculators and is challenged by rapidly appearing variants of the virus [12, 13]. Additionally, there is a large group of people, e.g. pregnant women, young children, people having certain comorbidities, who are not recommended to become vaccinated due to their medical condition [14]. With respect to the repurposing of drugs and the development of new agents, their efficiency is far from satisfactory [15]. Therefore, the discovery of novel anti-viral drugs specific to SARS-CoV-2 is urgently required.

The CoV genome is composed of a single, 5′-capped and 3′-polyadenylated RNA [16]. The first two-thirds of the genome encode two overlapping open reading frames (ORFs), ORF1a and ORF1b that produce polyproteins 1a and 1ab in a ribosomal frameshift. Downstream of this region, the remaining one-third of the genome encodes structural proteins (spike, envelope, membrane, and nucleocapsid) and several ORFs that produce the accessory proteins [17]. The polyproteins 1a and 1ab are then processed post-translationally by a virus-encoded protease, known as non-structural protein 5 (nsp5 or main protease) to produce other non-structural proteins (nsps) [18].

Among these proteins non-structural protein 1 (nsp1) encoded by the gene closest to the 5′ end of the viral genome received extensive attention. Nsp1 proteins of β-CoVs display similar biological functions in suppressing host gene expression [19, 20] and inhibiting the innate immune response to virus infection [18, 19]. SARS-CoV and SARS-CoV-2 nsp1 induce a near-complete shutdown of host protein translation by a three-pronged strategy: First, it binds to the small ribosomal subunit and stalls canonical mRNA translation at various stages [21, 22]. Second, nsp1 binding to the host ribosome leads to endonucleolytic cleavage and subsequent degradation of host mRNAs [21]. The interaction between nsp1 and a conserved region in the 5′ untranslated region (UTR) of viral mRNA prevents the shutdown of viral protein expression as a self-protection mechanism [23, 24]. Third, nsp1 interacts with heterodimeric host messenger RNA (mRNA) export receptor NXF1-NXT1, which is responsible for nuclear export of cellular mRNAs, causing a significant number of cellular mRNAs to be retained in the nucleus during infection [25]. Likewise, MERS-CoV nsp1 also exhibits a conserved function to negatively regulate host gene expression, despite employing a different strategy that selectively inhibits the host translationally competent mRNAs [26]. After infection, nsp1 also contributes to the evasion of the innate anti-viral immune response, mainly by antagonising interferon (IFN) induction and downstream signalling. In view of the crucial roles of nsp1 in the replication and virulence of CoVs, it represents an attractive target for both vaccine development and drug discovery.

The crystal structure of the N-terminal domain of SARS-CoV-2 nsp1 has been determined to 1.65 Å resolution (PDB entry 7K3N) [27] and the small C-terminal domain in complex with the human 40S ribosome (PDB entry 7K5I) [24] has been determined by cryo-electron microscopy providing useful structural information although the entire nsp1 structure could not be determined, probably due to its flexible linker region connecting these two distinct domains.

In this project we optimised crystallisation conditions and determined the structure of the N-terminal domain of nsp1 to atomic resolution of 0.95 Å, which served as a basis for fragment-based screening via x-ray crystallography. We identified two ligand binding pockets on nsp1, which bind a range of fragments. The fragment hits were further characterised by orthogonal biophysical assays. These fragments will serve as a basis for structure-based drug design.

## Results and discussion

### Structure determination of nsp1 to atomic resolution

SARS-CoV-2 is a protein of 180 residues consisting of an N-terminal domain (residues 1 to 128), followed by a linker region (residues 129 to 148) and a smaller C-terminal domain (residues 149 to 180). We generated an expression construct covering almost the entire globular domain (residues 10 to 126) (Figure 1A). A DNA insert coding for residues 10 to 126 of SARS-CoV-2 nsp1, named nsp1_10-126_ throughout the manuscript, was codon optimised for expression in *E. coli* and the protein was expressed and purified. Nsp1_10-126_ was subjected to crystallisation trials using published crystallisation conditions[27], but this did not yield any crystals. We therefore tested a variety of commercial crystallisation screens, which yielded three types of crystals with distinct morphology (supplementary material section Table S1). Crystals were cryo-cooled in liquid nitrogen, tested for diffraction quality and data were processed, all yielding the same space group P4_3_2_1_2 with very similar cell parameters (*a*=*b*=36.81 Å, *c*=140.96 Å, α=β=γ=90°) and identical to the published space group[27].

**Figure 1.**
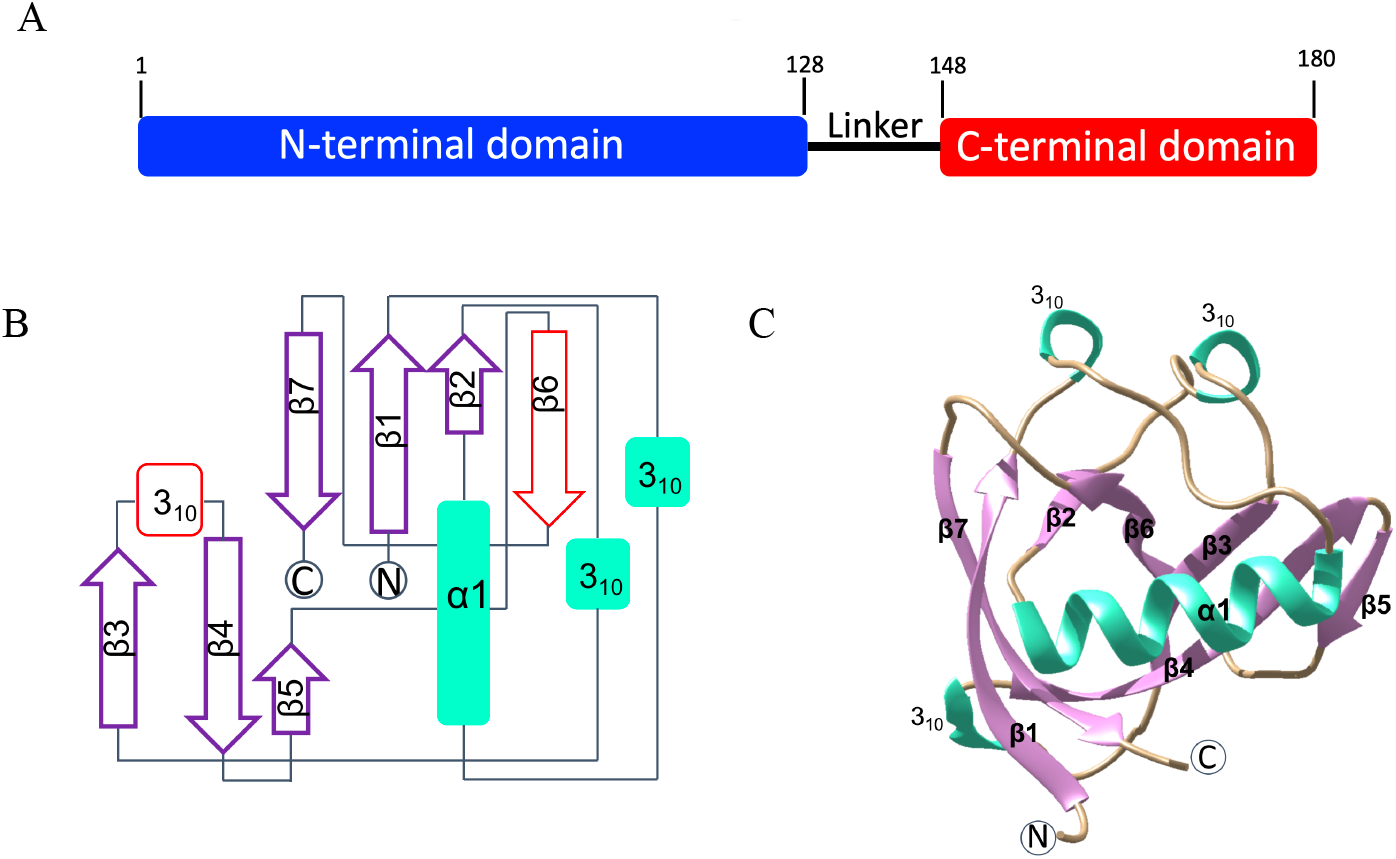
The structure of SARS-CoV-2 nsp1_10-126_. A) Bardiagram of the nsp1 domain arrangement, including the N-terminal domain (blue), the flexible linker region (black) and the C-terminal domain (red). B) Topological arrangement of SARS-CoV-2 nsp1_10-126_ at high resolution, where newly identified structural features are coloured in red. C) Cartoon representation of the structure. The secondary structure elements are depicted in different colours in the right panel with α-helixes coloured in green, β-strands shaded in purple, and loops are shown in tan.

The structure of Nsp1_10-126_ was determined by molecular replacement (MR) and refined to atomic resolution at 0.95 Å. Data collection and refinement statistics are summarised in Table 1. The asymmetric unit contains one molecule of nsp1. The model covers the sequence from Glu10 to Asn126. Currently, it is the highest resolution structure of a CoV Nsp1.

**Table 1.** Data collection, structure determination and refinement statistics for SARS-CoV-2 nsp1_10-126_. Data in parenthesis correspond to the highest resolution shell.

|  | SARS-CoV-2 nsp1<sub>10-126</sub> |
| --- | --- |
| Wavelength [Å] | 0.82656 |
| Resolution range [Å] | 32.63 - 0.95 (0.984 - 0.95) |
| Space group | P4<sub>3</sub>2<sub>1</sub>2 |
| Unit cell parameters [Å, °] | 36.81, 36.81, 140.96, 90, 90, 90 |
| Molecules per asymmetric unit | 1 |
| Total reflections | 469243 (66169) |
| Unique reflections | 62200 (9745) |
| Multiplicity | 7.5 (6.8) |
| Completeness [%] | 99.4 (98.2) |
| Mean I/sigma(I) | 11.31 (0.51) |
| Wilson B-factor (Å<sup>2</sup>) | 12.1 |
| R<sub>meas</sub> [%] | 6.4 (309.0) |
| CC<sub>1/2</sub> | 0.998 (0.172) |
| Reflections used in refinement | 61563 (5572) |
| R<sub>cryst</sub> / R<sub>free</sub> [%] | 16.9 (36.1) / 18.8 (38.1) |
| Total no. of non-hydrogen atoms (protein) | 1088 |
| No. of protein/ligand/solvent residues | 117 / 0 / 140 |
| RMSD bonds length, bond angles [Å, °] | 0.012 / 1.44 |
| Ramachandran<br>favoured/allowed/outliers/rotamer outliers [%] | 97.4 / 2.6 / 0/0 / 0/0 |
| Clashscore | 8.3 |
| Average B-factor/protein/ligands/solvent | 23.4 / 21.2 / 38.5 |

The structure of SARS-CoV-2 nsp1_10-126_ features a unique topological arrangement resulting in the formation of a seven-stranded (n = 7) β-barrel that is primarily antiparallel, except for strands β1 (His13 – Val20) and β2 (Cys51 – Val54) (Figures 1B and 1C). Additional major structural features include helix α1 (Val35 – Asp48), which is positioned as a cap along one opening of the β-barrel, three 3_10_ helices that run parallel to each other, and the strand β5 (ILe95 – Tyr97), which is not part of the β-barrel but forms a β-sheet interaction with the β4 strand (Val84 – Leu92). Compared to the recently published structures determined at 1.6 Å and 1.77 Å resolution (PDB entries 7K3N and 7K7P) [27] [28], our atomic structure identified a third 3_10_ helix in the flexible loop region between β3 and β4 (residues 80 to 82). As the density indicates that this (loop) region may have various conformations, we tentatively fitted one conformation into the density. In addition, our structure also reveals a new strand, β6, and shows a longer strand β5 than the previously published structures. This more complete model refined to higher resolution provides a more accurate structural picture of nsp1 and will serve as an excellent starting point for fragment-based screening.

### Fragment screening via X-ray crystallography

Fragment screening was performed by soaking 584 fragments from the Maybridge Ro3 library into nsp1_10-126_ crystals followed by validation through X-ray diffraction experiments in fully automated mode at beamline MASSIF-1 [29, 30]. In addition, 40 nsp1 crystals without fragments were prepared to construct the Ground State (GS) model for data analysis in the multi-crystal software system PanDDA [31]. The overall resolution of data sets for the GS was 1.3 Å. 398 fragments were accepted for PanDDA analysis. Datasets that were rejected for the high R_free_ values (14) were reprocessed and manually inspected. Potential fragment hits were discovered by PanDDA (34), and nine of them were verified in the single crystal system by manual inspection in COOT [32] followed by refinement in Phenix [33]. These fragment hits can be defined in two groups binding to two distinct ligand binding sites named binding pocket 1 and binding pocket 2. Here we characterise five selected fragment hits. Data collection and refinement statistics for SARS-CoV-2 nsp1_10-126_-fragment complexes are summarized in Table S2 of the supplementary material section and the chemical structures of the fragment hits are shown in Figure 2. Fragments **10B6, 11C6** and **5E11** represent hits in which a phenyl group is connected to a 5-membered heterocyclic ring system. The second group of fragments represented by **7H2** and **8E6** contain a single phenyl ring system containing single or double substitutions.

**Figure 2.**
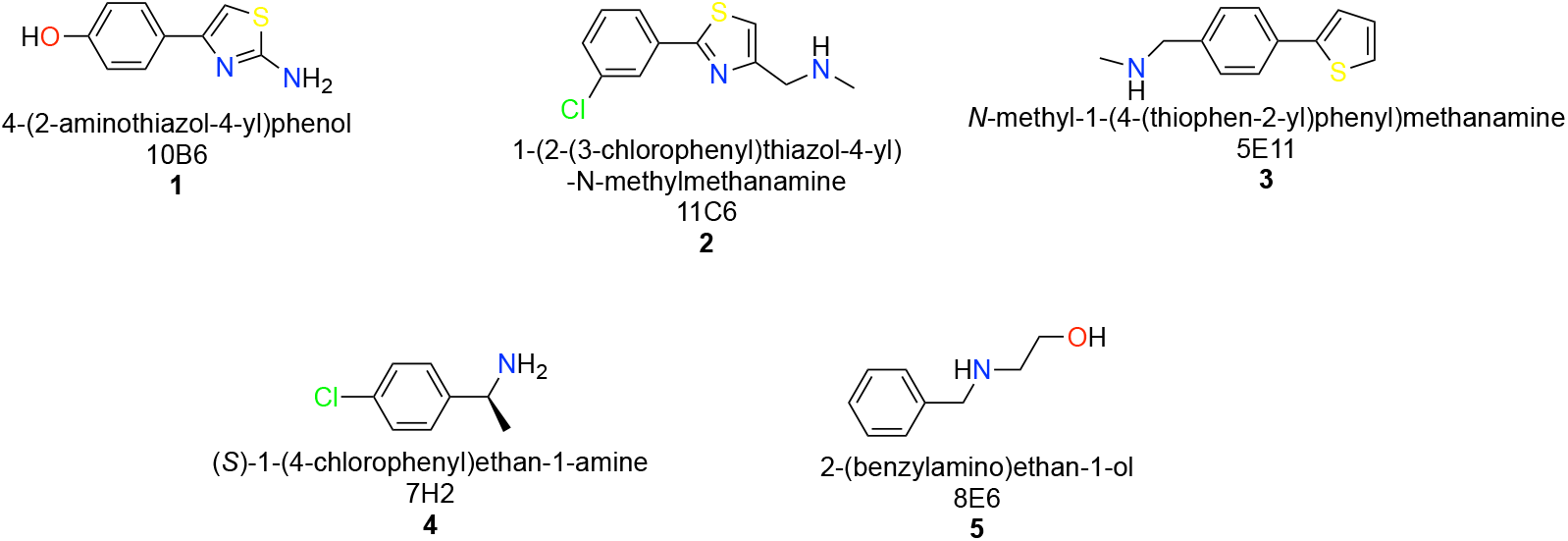
Chemical structures, systematic names, plate positions and numbering of fragment hits binding to two distinct binding sites in SARS-CoV-2 nsp1_10-126_.

### Identification of ligand binding sites in SARS-CoV-2 nsp1

By overlaying all nsp1-fragment complexes, we could identify two distinct ligand-binding pockets. The majority of fragment hits, 3, bind to ligand binding site I, whereas 2 of the 5 hits bind to the shallower binding site II (Figure 3a). Ligand binding site I involves the N- and C-termini, strands β1 and β7 as well as residues of helix α1. Ligand-binding site II is formed by nsp1 and one of its symmetry mates involving the loop region between the first 3_10_ helix and helix α1, two residues of α1 and residues in the loop connecting strands β6 and β7. Residues from the symmetry mate involved in binding are located in and close to the first 3_10_ helix and in the second 3_10_ helix. The shortest distance between the two binding pockets is about 18 Å and seems to be challenging to bridge for fragment linking strategies. The characteristics of these two pockets are summarised in Table 2.

**Table 2.**
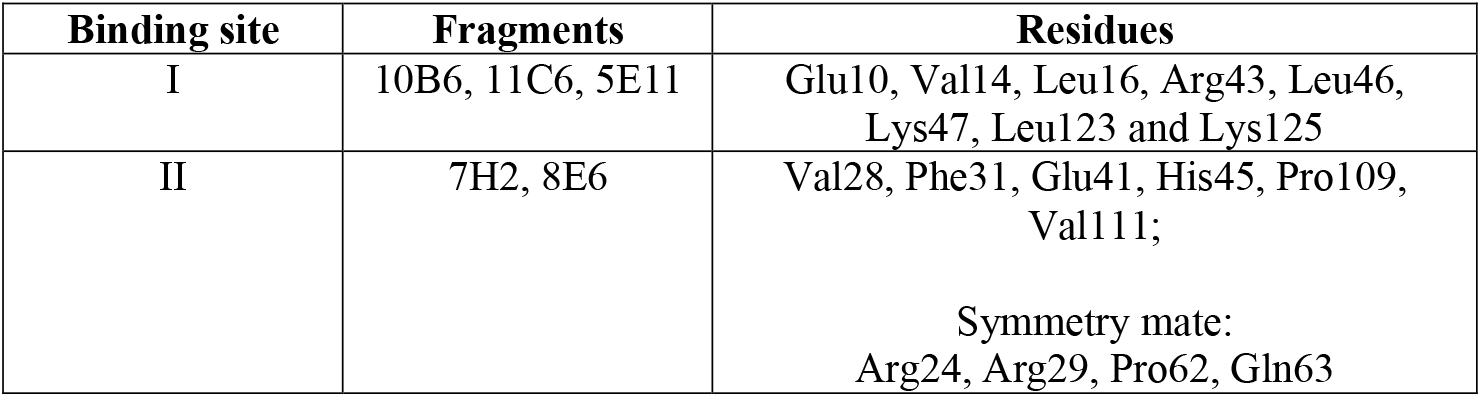
List of SARS-CoV-2 nsp1_10-126_ amino acids defining binding sites I and II.

**Figure 3.**
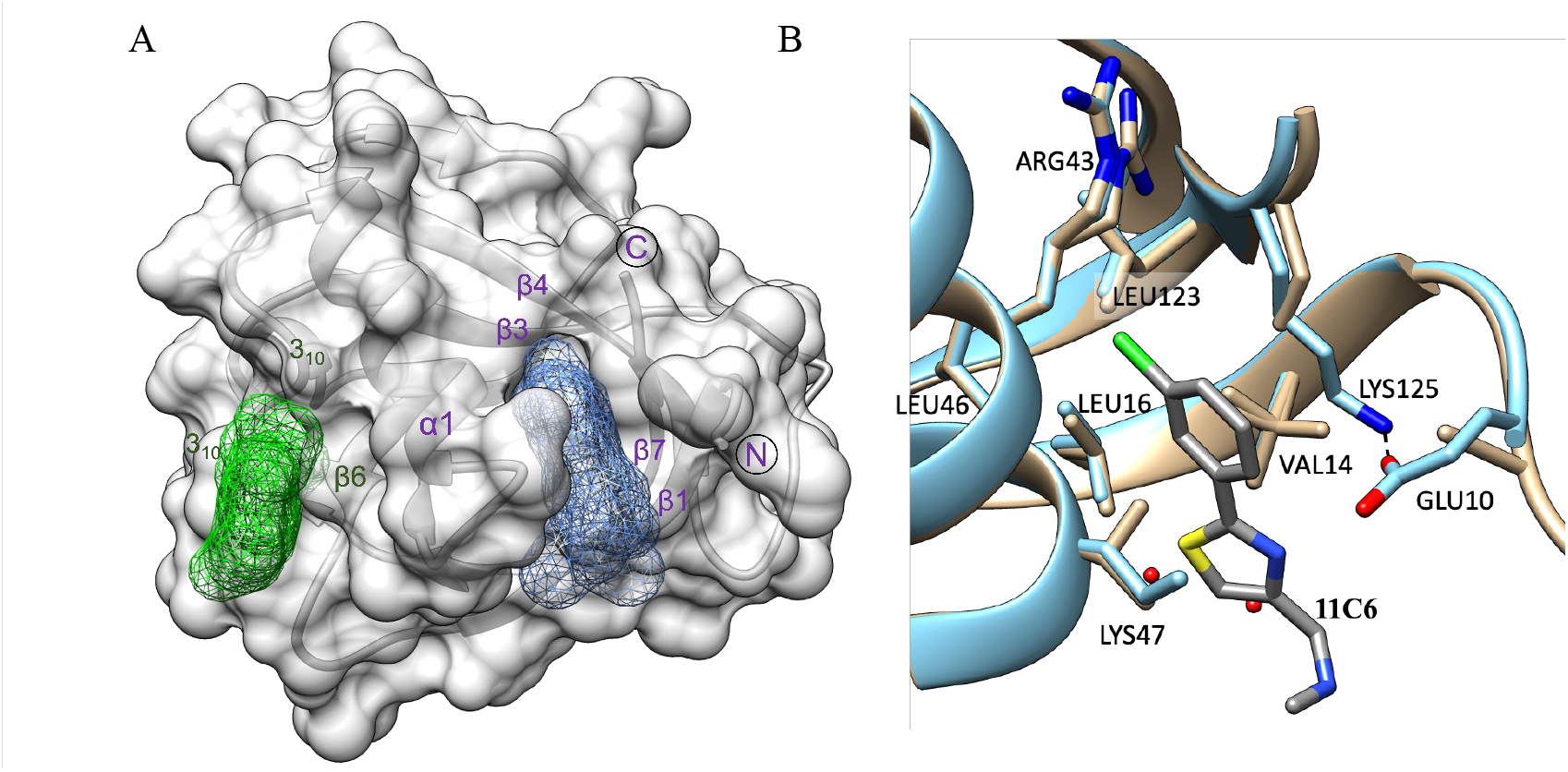
Binding sites I and II in SARS-CoV-2 nsp1_10-126_. A) Surface representation of SARS-CoV-2 nsp1_10-126_ shown in light grey indicating binding sites I (blue) and II (green). The secondary structures forming the two binding pockets are labelled in the corresponding colours with α1 contributing to the formation of both pockets. B) Overlay between native nsp1 (beige) and the SARS-CoV-2 nsp1_10-126_-11C6 complex (blue) showing the structural changes induced by fragment binding.

### Fragment hits binding to pocket I induce small structural rearrangements

Having high-resolution native and fragment-bound structures allows us to discuss ligand-induced structural changes in detail. Fragment **11C6** was chosen as an example. With respect to binding site I, Glu10, Lys47 and Lys125 show the most obvious changes. Whereas in the native atomic structure the side chain of Lys125 is only partially visible, it is fully visible in the ligand bound structure, as it runs parallel to the fragment and forms hydrophobic interactions with one of the phenol rings and displays a hydrogen bond interaction with Glu10 (Figure 3b). In the fragment bound structure the Glu10 side chain density is therefore mostly visible, while it is absent in the native structure. The density for the Lys47 side chain is also only partially visible in both native and fragment bound structures but some positive density in the native structure suggests one conformation extending toward the space, which the fragment would occupy. In the native structure, two water molecules are located in, or close to, the space otherwise occupied by the 3-chlorophenyl group of the ligand, but these are absent in the fragment bound structure. In conclusion, binding of fragments to this site leads to a stabilisation of surrounding residues indicating a small but noticeable induced fit mechanism caused by fragment binding. In contrast, binding of fragment **7H2** to binding site II does not induce any significant structural changes in its surrounding environment (data not shown).

### Detailed description of SARS-CoV-2 nsp1_10-126_-fragment interactions

Fragments hits **10B6, 11C6** and **5E11** are chemically related as they contain a substituted phenyl ring system connected to a five-membered heterocyclic thiophen or thiazol ring system. All three bind to ligand binding site I, establishing various interactions.

**10B6** establishes various hydrogen bond interactions with a range of residues of binding pocket I (Figure 4a). The 2-amino group establishes a hydrogen bond interaction with a water molecule. Fragment **11C6** maintains the overall binding conformation in the same pocket (Figure 4b). However, this fragment contains a chloride substituent in the *meta*-position instead of the *para*-substituted hydroxyl group. Binding occurs exclusively through hydrophobic interactions, but compared to **10B6, 11C6** displays significantly less interaction with nsp1. Interestingly, the orientation for fragment **5E11** is reversed whereby the 6- and 5-membered rings switch positions (Figure 4c). Overall, many hydrophobic interactions of the fragment hit with residues in binding pocket I are maintained, but the nitrogen of the Lys47 side chain now establishes a hydrogen bond interaction with the sulphur of the thiophen ring system, while a N-methyl methanamine substituent on the 6-membered ring system does not establish a hydrogen bond interaction. This example shows the importance of high-quality and high-resolution structural information as mostly flat fragments can be easily fit into the electron density in various orientations and a wrong orientation of the ligand in subsequent SAR studies could easily lead to incorrect conclusions.

**Figure 4.**
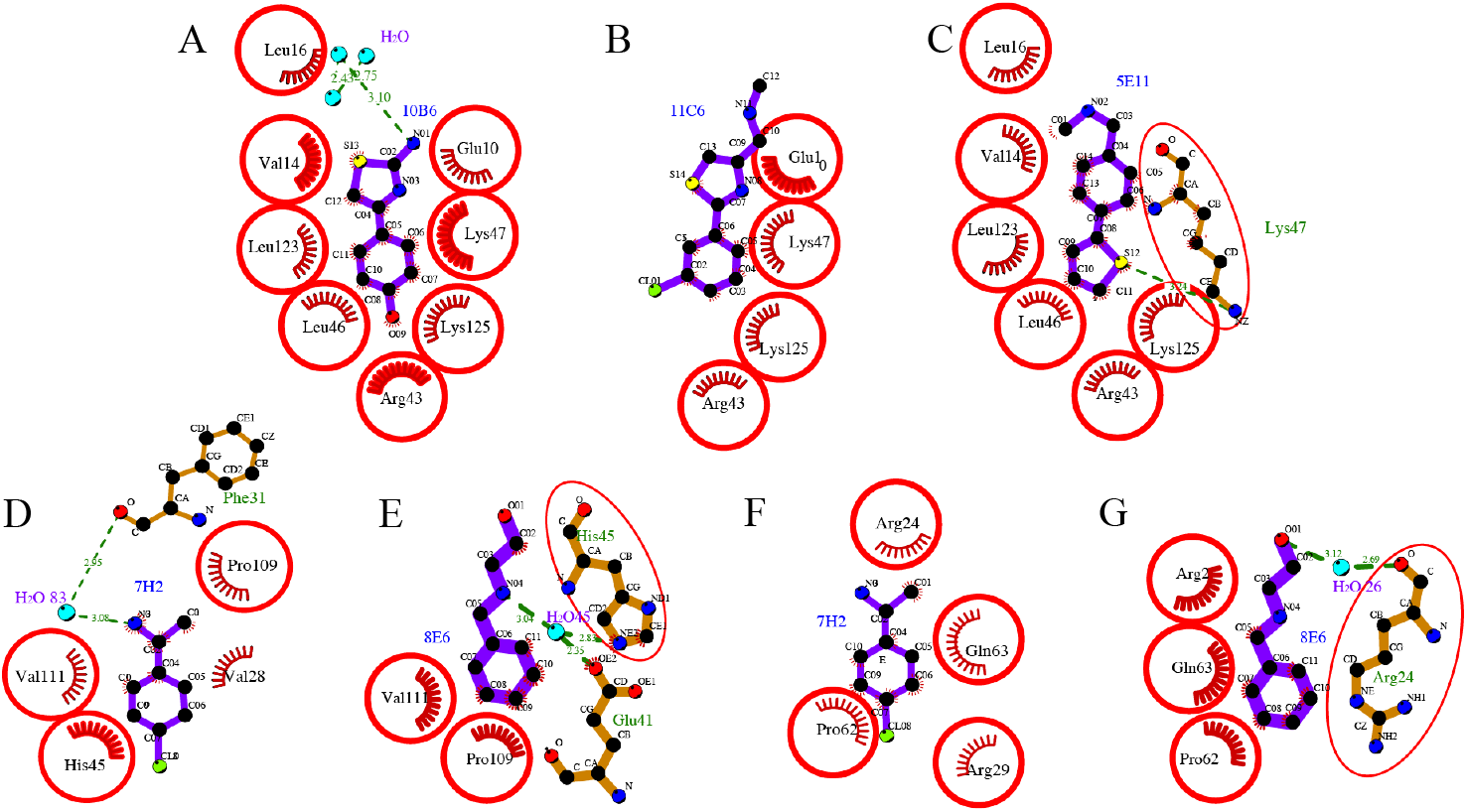
Molecular interactions of fragment hits binding to SARS-CoV-2 nsp1_10-126_. Hydrophobic interactions are displayed by red half moons and identical residues are shown with a red circle. Hydrogen-bond interactions are shown with a dotted green line. Three fragment hits (A) **10B6**, (B) **11C6** and (C) **5E11** interacting with the residues of SARS-CoV-2 nsp1_10-126_ in binding pocket I. The binding conformation is switched in **5E11**, in which the 5- and 6-membered ring system exchange their positions compared to **10B6** and **11C6**. Fragment hits **7H2** and **8E6** interacting with the residues located in binding site II. **7H2** binding to D) SARS-CoV-2 nsp1_10-126_ and F) a symmetry mate. **8E6** interacting with E) SARS-CoV-2 nsp1_10-126_ and G) a symmetry-related molecule. The substituent of **8E6** is not visible in the electron density.

Fragments **7H2** and **8E6** bind to a distinct site on SARS-CoV-2 nsp1_10-126_ named binding pocket II (Figure 4 d-g). The common feature of these two hits is that they both possess a single substituted phenyl ring system.

**7H2** displays various hydrophobic interactions with residues of SARS-CoV-2 nsp1_10-126_ (Figure 4d). The ethan-1-amine group establishes a hydrogen-bond interaction through a water molecule with the main chain oxygen of Phe31. The opposite chloride substituent does not seem to contribute to binding to SARS-CoV-2 nsp1_10-126_. It is noticeable that compared to fragments binding to pocket I, **7H2** has a reduced number of interactions with residues of the protein, probably because this ligand binding site II is shallower than pocket I. Compared to **7H2, 8E6** is a mono-substituted fragment lacking the chloride substituent but carrying an amin-1-ethan-1-ol substituent instead of the ethan1-1-amine (Figure 4e). As the data for this complex were particularly good, with a resolution of 1.18 Å, we placed the fragment in all possible orientations into the electron density and conducted refinement for six distinct orientations. The best refinement was obtained in which the substituent points towards the solvent, establishing a hydrogen-bond interaction with the side chains of Glu41 and His45 through a water molecule. Despite essentially observing electron density for only part of the fragment, **8E6** also establishes hydrophobic interactions with Pro109 and Val111, similar to **7H2**. In contrast to **10B6, 11C6** and **5E1, 7H2** (Figure 4f) and **8E6** (Figure 4g) also establish interactions with a symmetry mate generating a set of novel interactions, which requires orthogonal assays to establish if there is a “real” binding site, or if these only bind in the presence of the arrangement dictated by the crystal.

### Orthogonal biophysical assays to further validate fragment hits

To further validate SARS-CoV-2 nsp1_10-126_-targeting fragment hits, we employed two orthogonal biophysical assays, thermal shift assays (TSA) and microscale thermophoresis (MST). For MST experiments, the binding affinities between SARS-CoV-2 nsp1_10-126_ and fragment hits were evaluated using the Monolith NT. 115 instrument with the pico-red fluorescent channel. The MST results of the fragments are summarised in Table 3. The K_d_ values vary between 475 μM and larger than 20 mM. Interestingly, fragment hits binding in the “deeper” binding site systematically show better K_d_ values than the two hits binding in the shallower site II which display weaker K_d_ values. Crystallographic analysis of the binding site for **7H2** and **8E6** revealed that a symmetry mate contributed to the binding of the two fragments to nsp1, raising the possibility that both proteins in the particular crystallographic arrangement are required for binding and that these two fragments may not bind to nsp1 in solution. However, the calculated K_d_ values clearly show that binding to a SARS-CoV-2 nsp1_10-126_ monomer in solution is sufficient.

**Table 3.** Summary of x-ray crystallography, MST and TSA results for fragment hits targeting SARS-CoV-2 nsp1_10-126_ and selected calculated properties. The **Δ**T_i_ values of SARS-CoV-2 nsp1_10-126_ in the presence of fragments obtained from nanoscale differential scanning fluorimetry (nanoDSF) and fragment affinities for SARS-CoV-2 nsp1_10-126_ determined by MST are shown. TSA and MST experiments were conducted at least in triplicate. The T_i_ of SARS-CoV-2 nsp1_10-126_ in the presence of 2.0% DMSO is 54.05°C, which was used as a control in ΔT_i_ calculations. The following chemical properties were calculated by Molsoft’s ICM Pro software[35]. MW, molecular weight; PSA, polar surface area; clogP, calculated logP; HBA, hydrogen bond acceptor; HBD, hydrogen bond donor; clogS, calculated solubility. *APUC: atypical protein unfolding curve.

| Fragment ID | Binding site | MST $K_d$ [mM] | TSA $T_i$ *± SD | TSA $\Delta T_i$ [°C] | TSA $\vert \Delta T_i\vert $-3SD [°C] | MW [Da] | MolLogP (mol) | HBA & HBD [No] | mol PSA [$\text{\AA}^2$] | MolLogS [Log(moles/L)] |
| --- | --- | --- | --- | --- | --- | --- | --- | --- | --- | --- |
| **5E11** | I | 0.48 ± 0.30 | APUC | / | / | 203.1 | 2.75 | 1, 1 | 13.0 | -3.526 |
| **10B6** | I | 0.56 ± 0.19 | 51.27 ± 0.15 | -2.51 | 2.06 | 192.0 | 2.48 | 2, 3 | 48.4 | -3.495 |
| **11C6** | I | 1.16 ± 0.22 | 53.78 ± 0.05 | -0.90 | 0.76 | 238.0 | 2.73 | 2, 1 | 22.4 | -4.518 |
| **7H2** | II | >20 | 53.55 ± 0.25 | 2.28 | 1.53 | 155.1 | 1.23 | 1, 2 | 20.6 | -2.565 |
| **8E6** | II | 8.32 ± 3.32 | 52.72 ± 0.22 | -0.83 | 0.16 | 151.1 | 0.51 | 2, 2 | 28.8 | -0.603 |

For TSA, nsp1 in the presence of an equal concentration of DMSO was used as a control to offset the influence of DMSO on the protein. The averaged change of inflection temperature (△T_i_) of the protein in the presence of fragments higher than three-fold of standard deviation (|ΔTi|-3*SD>0) was regarded as statistically significant. As can be seen from Table 3, except for **5E11**, which showed atypical unfolding curves, all fragment hits influenced the unfolding property of SARS-CoV-2 nsp1_10-126_. A stabilisation effect was observed for **7H2**, while destabilisation effects were measured for **8E6, 10B6** and **11C6**. In this regard, TSA may not be suitable yet for the validation of hits against SARS-CoV-2 nsp1_10-126_, as we have recently also observed for another SARS-CoV-2 protein, nsp10 [34]. With respect to physicochemical properties, calculated values for fragments are well within the expected range, indicating good starting points for structure-based drug design.

### Cross inhibition of SARS-CoV-2 nsp1_10-126_ fragment hits on SARS and MERS homologues

To expand our investigation to other β-coronaviruses of medical relevance, fragment hits identified by x-ray crystallography were also tested on SARS-CoV-1 and MERS nsp1s to investigate potential cross-inhibitory effects using MST. With the high sequence identity of 86.3% between SARS-CoV-2 and SARS-CoV-1 nsp1s, only 7H2 shows no binding to SARS-CoV-1 nsp1 whereas **11C6** showed strong cross-inhibitory effects. With lower sequence identity between MERS and SARS-CoV-2 nsp1, only **10B6** showed moderate binding affinity to MERS nsp1 (Table 4). This may be a first indication that developing pan inhibitors across medically relevant β-coronavirus by targeting nsp1 is promising.

**Table 4.** Cross-inhibitory effects of fragment hits identified against SARS-CoV-2 nsp1_10-126_ and their estimated inhibitory effects on the nsp1 homologues from SARS-CoV-1 and MERS. *N.i, no inhibition

| Fragment hit | SARS-CoV-2 nsp1<br>MST, $K_d$ [mM] | SARS-CoV-1 nsp1<br>MST, $K_d$ [mM] | MERS nsp1<br>MST, $K_d$ [mM] |
| --- | --- | --- | --- |
| **5E11** | $0.48 \pm 0.30$ | $12.76 \pm 24.13$ | >20 |
| **10B6** | $0.56 \pm 0.19$ | >20 | $4.92 \pm 4.19$ |
| **11C6** | $1.16 \pm 0.22$ | $0.38 \pm 0.14$ | n.i. |
| **7H2** | >20 | n.i. | n.i |
| **8E6** | $8.32 \pm 3.32$ | >20 | >20 |

### The two novel binding sites in SARS-CoV-2 nsp1_10-126_ are not conserved among SARS-CoV-1 nsp1 and MERS nsp1

To probe the homology of SARS-CoV-2, SARS-CoV-1, and MERS nsp1s, the sequences of the N-terminal domain of the three nsp1s were compared using structure-based sequence alignment (Figure 5). Whereas nsp1 is highly conserved between SARS-CoV-1 and SARS-CoV-2 (86.3% identical residues, 8.6% strongly similar, 0.9% weakly similar, 4.3% different), the protein sequence from MERS differs significantly (19.4% identical residues, 21.0% strongly similar, 11.3% weakly similar, 48.4% different). A major difference between MERS and the two SARS sequences is the presence of four insertions in loop regions of the protein. Consequently, most residues forming part of the two ligand binding pockets for the five fragment hits are therefore conserved between SARS-CoV-2 nsp1 and SARS-CoV-1 nsp1 but differ in MERS nsp1, in particular in the binding site II.

**Figure 5.**
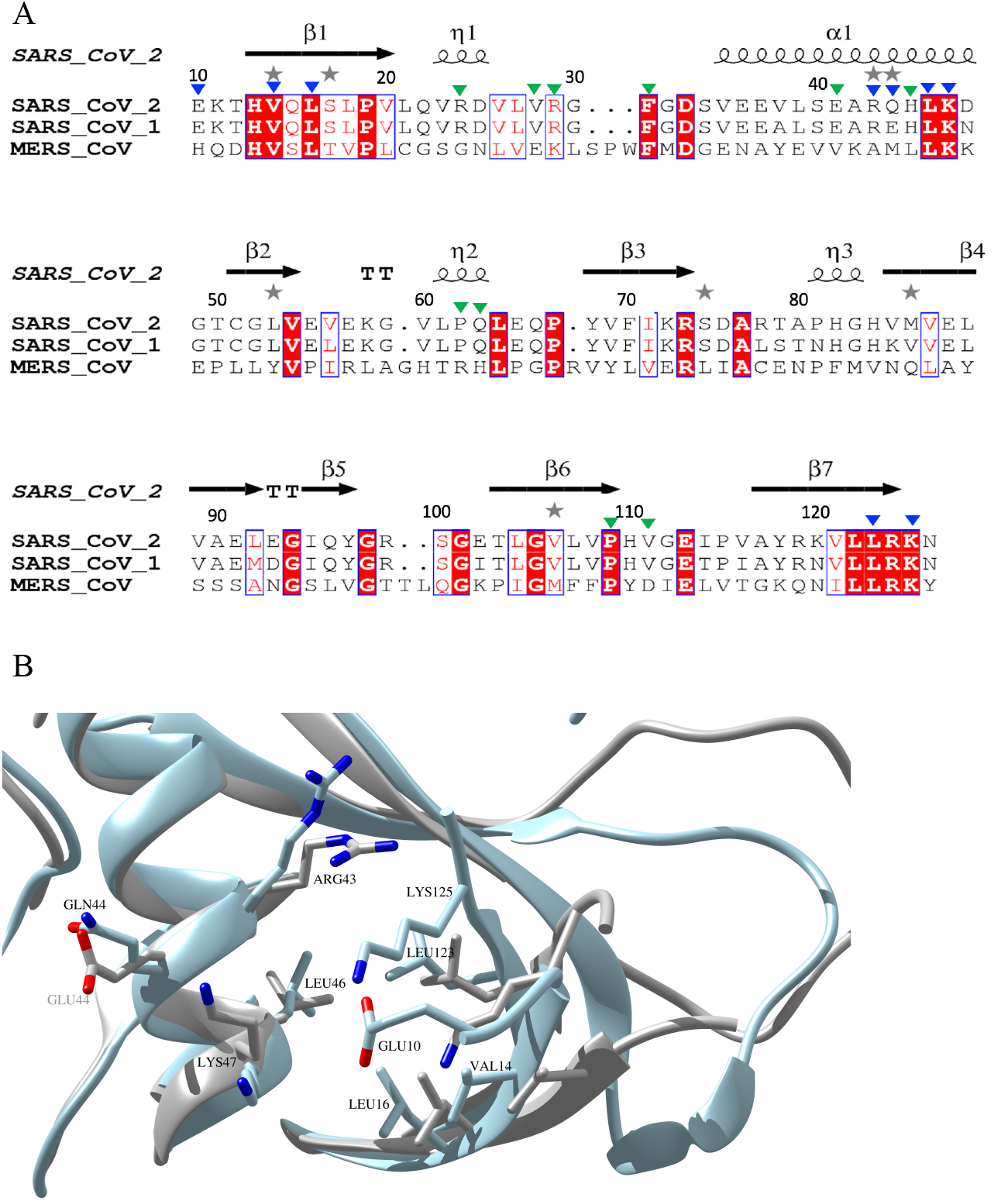
Sequence and structural alignment of the N-terminal domain of three medically relevant coronaviruses SARS-CoV-2, SARS-CoV-1 and MERS nsp1s. A) Structural protein sequence alignment of SARS-CoV-1, SARS-CoV-2 and MERS. Key residues in binding pockets I and II are indicated by blue and green triangles, respectively. The sequences were aligned using CLUSTAL W multiple alignment [36] and the secondary structure elements were extracted using ESPript 3.0 [37]. B) Overlay of SARS-CoV-1 (coloured in light grey) and SARS-CoV-2 (coloured in light blue) highlighting the structural differences around binding pocket I. Pairwise key residues of the two nsp1s in pocket I interacting with fragment hits are shown in sticks. Glu10 is absent in the NMR structure of SARS-CoV-1 nsp1 (PDB entry 2HSX).

As the structure of MERS nsp1 has not yet been determined, the structures of the two available SARS-CoV-1 and SARS-CoV-2 nsp1 proteins were overlaid highlighting key residues in the two binding pockets. For binding pocket I, all key residues of SARS-CoV-2 nsp1_10-126_ involved in fragment binding are conserved in N-terminal domain of SARS-CoV-1 nsp1. However, due to the noticeable displacement of the secondary structural elements in this region between the two proteins, these conserved residues are oriented in different directions or located at different locations (Figure 5b). As even minor changes in these residues could greatly influence the interaction with fragment hits, these structural changes could probably alter the binding for fragment hits on N-terminal domain of SARS-CoV-1 nsp1, explaining their reduced binding affinity compared to SARS-CoV-2 nsp1. In binding pocket II, structural changes are less pronounced, but may be sufficient to abolish weak binding affinities of the fragments. Based on these structural comparisons, we therefore hypothesise that although SARS-CoV-1 nsp1 and SARS-CoV-2 nsp_10-126_ share high sequence identities in binding sites I and II, the structural differences in these pockets between SARS-CoV-2 nsp_10-126_ and N-terminal domain of SARS-CoV-1 nsp1 probably explain the loss of binding affinity of fragment hits **5E11, 10B6, 7H2**, and **8E6** on N-terminal domain of SARS-CoV-1 nsp1. A full understanding would require obtaining structural information on MERS and SARS fragment complexes.

### The predicted RNA binding and validated DNA polymerase *α*–primase binding regions are in close proximity to fragment binding sites I and II

The N-terminal nsp1 domain has been reported to release the expression shutoff for viral RNA by direct interaction with the stem loop 1 (SL1) [38]. Therefore, the spatial relationship between the binding pockets and the interaction regions was first investigated for potential overlaps of key residues for both fragment hits and mRNA binding. According to computational docking experiments between nsp1 and the host RNA SL1 region, nsp1 displays direct interactions mainly through residues 11-17, 118-130, and 144-148, where hydrogen bonds and ionic interactions were observed for a range of key residues [38]. Among these, Lys125 is both an important contributor to the two types of interactions with RNA but also interacts with fragment hits in binding pocket I (Figure 6A). In view of the close proximity and this shared key residue, we speculate that targeting binding pocket I could interfere with RNA binding to nsp1, and subsequently impact on viral RNA expression.

**Figure 6.**
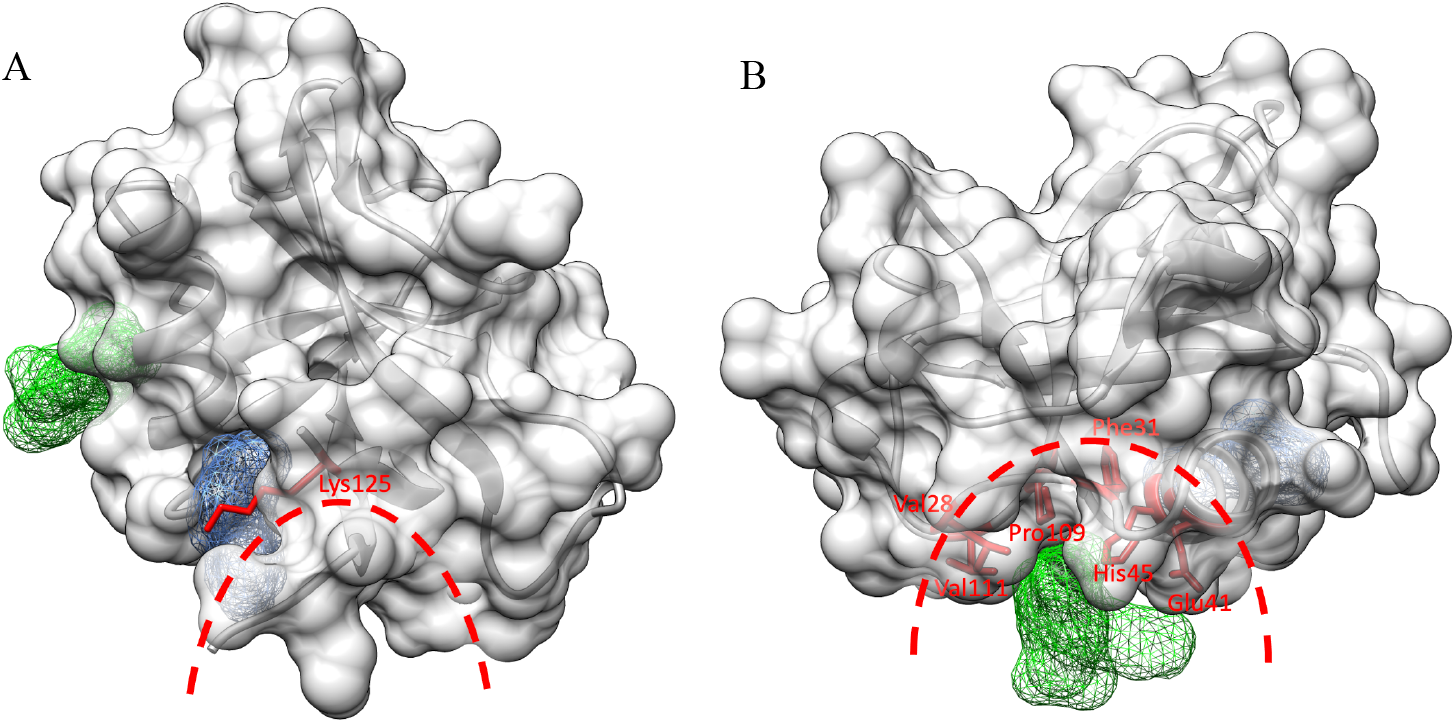
The close relationship in space between two binding pockets and targets in host cells. A) The close spatial proximity between binding pocket I (blue mesh) and the predicted RNA binding region, indicated by the red dashed line. The shared key residue Lys125 is shown in red as a ball-and-stick model. (B) The spatial overlap between binding pocket II (green mesh) and the interacting surface with the primosome is indicated by the red dashed line. The shared key residues Val28, Phe31, Glu41, His45, Pro109 and Val111 are shown in red as ball-and-stick models.

The primosome, the complex of DNA polymerase α (Pol α) and primase that is responsible for initiating DNA synthesis during DNA replication has recently been reported to be an important human host protein targeted for SARS-CoV-2 nsp1 [39, 40]. In addition, the observation of disease-associated mis-splicing of POLA1 coding for the catalytic subunit of Pol α resulting in abnormal IFN I response and auto-inflammatory manifestation reveals the participation of Pol α in innate immunity [41, 42]. By inspecting the interaction surface between SARS-CoV-2 nsp1 and the catalytic subunit of Pol α from the determined cryo-electron microscopy structure of the SARS-CoV-2 nsp1-primosome complex [43], we found that the fragment binding pocket II is located at the centre of the interaction interface and shares several key residues (Val28, Phe31, Glu41, His45, Pro109 and Val111) with the primosome (Figure 6B). Therefore, we suggest that targeting binding pocket II could interrupt the interaction between SARS-CoV-2 nsp1 and the primosome, thus releasing the inhibition on primosome in DNA synthesis and immune response.

### Development of resistance through mutations in nsp1 ligand binding pockets?

Recently, Mou and co-workers, investigated close to 300,000 fully sequenced SARS-CoV-2 genomes for mutations in nsp1[44]. For full-length nsp1 containing 180 residues they identified 933 non-synonymous mutations in the entire coding region, identifying at least one mutation for each single residue. The 19 most common mutations with a frequency between 113 and 1122 were reported. By comparing residues forming binding pocket I (Table 2) with the 19 residues with the highest frequency we conclude that none of the residues of this binding site belongs to the group with highest frequency, indicating that these residues do not seem to be among the group of residues that mutate easily. However, all residues in binding site I display some tendency to mutate with frequencies between 4 and 9. As binding pocket II is very shallow, only a limited number of SARS-CoV-2 nsp_10-126_ residues establish interactions with the fragment hits. Interestingly, one of the three residues, His45, belongs to the group of residues with most frequent mutations. However, since the interaction occurs via the main chain but not through the side chain, this mutation should have no significant effect, in case they do not significantly alter the local structure. In contrast, in the symmetry mate, Arg24 is one of the residues forming the ligand binding pocket. This residue shows the tendency to mutate with the highest frequency (1122) of all residues. In conclusion, binding pocket II is not only shallower than binding pocket I, but its residues involved in the formation of the pocket also show a higher tendency to mutate. We hypothesise that SARS-CoV-2 may be able to overcome inhibitors by either mutating key residues involved in inhibitor binding, or through primary resistance as mutations may already be present in the virus.

### The rationale for targeting N- and C-terminal domains of nsp1

After infection of host cells, SARS-CoV-2 nsp1 can bind to the 40S subunit of host ribosomes to block the entry of the host mRNA expression channel through its C-terminal domain[45]. The C-terminal domain is connected by a linker region to its N-terminal globular domain, which remains tethered around the entry site [23]. As a result, host mRNAs are inaccessible to the expression cleft. Therefore, the host ribosome is hijacked by SARS-CoV-2 nsp1 to mainly serve the production of viral proteins. Targeting the C-terminal SARS-CoV-2 nsp1 domain would thus represent a valid strategy to interfere with its binding to the 40S subunit and closure of the channel in the small ribosome subunit. However, this approach remains challenging as this small domain appears to be unstructured [46], only folding into two helices upon binding to the ribosome [47].

Residues in the N-terminal and linker region of SARS-CoV-2 nsp1 are not involved in docking into the mRNA entry channel, but they do stabilise its association with the ribosome, enhancing its restriction of host gene expression [48]. Ribosome-bound SARS-CoV-2 nsp1 further induces endonucleolytic RNA cleavage in the 5’-UTR of host mRNAs [49, 50]. However, viral mRNA with a special stem-loop 1 (SL1) within the leader sequence in the 5’-UTR can circumvent the SARS-CoV-2 nsp1-mediated translational inhibition and cleavage by directly interacting with the N-terminal domain and the linker region of the protein, ensuring sufficient viral protein expression [38, 51, 52]. It was also reported that the activity of SARS-CoV-2 nsp1 in translation termination is provided by its N-terminal domain by binding with the 80S ribosomes translating host mRNAs and removing them from the pool of the active ribosomes [53]. Additionally, the interaction between the N-terminal globular domain of SARS-CoV-2 nsp1 and the primosome, which is essential for host DNA replication, identified by biochemical and structural characterisation, suggests that targeting the primosome is part of a novel mechanism driving SARS-CoV-2 infections [43]. In view of the important roles played by both N- and C-terminal domains in host gene expression and escape of viral mRNA in translational repression, nsp1 appears to be a potential drug target.

In conclusion, we identified two ligand-binding pockets in the N-terminal domain of nsp1 that accommodate fragments and described their detailed molecular interactions. Whereas the first ligand binding site can be considered as a “true” pocket establishing multiple interactions between residues of SARS-CoV-2 nsp_10-126_ and fragment hits, the second ligand binding pocket is shallower with a reduced number of protein-fragment interactions and weaker K_d_ estimates. In an accompanying paper, we have used extensive molecular dynamics simulations and fragment-based screening to highlight the cryptic nature of these pockets (**doi:** https://doi.org/10.1101/2022.05.20.492819). Furthermore, our fragment screening work is also an excellent example of how difficult it can be to fit fragment hits into positive electron density, despite high resolution between 1.1 to 1.3 Å and excellent data quality, when fragment hits are predominantly flat, substituents around the core chemical structure are flexible and fragment hits also interact with symmetry mates in the crystal. MST assays conducted using full-length nsp1 from SARS and MERS, two previously known β-CoVs with severe medical impact, to investigate cross-inhibitory effects provide a first hint of how challenging it may be to develop pan CoV inhibitors against diverging medically relevant CoV such as SARS-CoV-2, SARS and MERS. The experimental identification of binding pockets and induced fit changes will be a good starting point for computational docking and screening. Nevertheless, fragment hits will act as templates in future analogue design, combined with structure-based optimisation. Fragment hits reveal druggable binding pockets on SARS-CoV-2 nsp_10-126_, which will provide novel insights into further drug design, but we assume that additional ligand binding pockets on SARS-CoV-2 nsp1 may be discovered in the future. Currently, the existence of two pockets allows for fragment growing and merging techniques but not fragment linking.

### Experimental

#### Subcloning, expression and purification of SARS-CoV-2 nsp1 constructs

Codon optimised DNA for expression in *E. coli* for the N-terminal domain of SARS-CoV-2 nsp1 was obtained by extracting the protein sequence of the N-terminal domain from PDB entry 7K7P. In frame restriction sites for NcoI and XhoI were added at the 5’ and 3’ and the synthesised insert was subcloned into expression vector ppSUMO-2, containing a Ulp1-cleavable N-terminal histag and SUMO domain, using the same restriction sites. The expressed nsp1 protein construct codes for residues 10 to 126 of SARS-CoV-2 nsp1 and includes two unspecific residues, Thr-Met, at the N-terminus of the protein due to the cloning strategy.

A similar cloning strategy was used to subclone the DNA coding for full-length SARS nsp1 (139 residues; NCBI reference sequence: NP_828860.2) and full-length MERS nsp1 (140 residues; NCBI reference sequence YP_009944292.1) into ppSUMO-2.

200 μL of competent *E. coli* BL21-Gold (DE3) cells (Agilent Technologies) was gently mixed with 1 μl ppSUMO-nsp1 expression plasmid and incubated on ice for 30 min. A heat shock at 42°C was employed for 45 sec followed by 2 min incubation on ice. Then, 400 μL of SOC medium (Sigma-Aldrich, UK) was added followed by incubation for 1 h on a shaker-incubator at 37°C, 500-600 rpm. 200 μL of transformed cells were spread evenly on LB agar (MP Biomedicals, LLC France) plates supplemented with 50 μg/mL kanamycin and incubated at 37°C overnight.

To express the SARS-CoV-2 nsp1_10-126_ construct, a single BL21-Gold (DE3) colony was picked and added into 35 mL of TB medium supplemented with 50 μg/mL kanamycin. The small culture was then incubated in a shaker at 37°C, 220 rpm overnight. Subsequently, the small culture was separately added into 12 Erlenmeyer flasks each containing 1 L of TB medium supplemented with 50 μg/mL kanamycin and incubated in a shaker at 37°C, 220 rpm for 5 h until the OD_600_ value reached on OD between 0.6 to 0.9. The large cultures were kept at 4°C for 20 min before being induced for protein expression using 0.1 mM (Isopropylthio-β-galactoside) IPTG and incubated at 18°C, 220 rpm for 24 h. The following day, the culture was harvested by centrifugation at 8000 g (Avanti^®^ J-E centrifuge from Beckman Coulter, rotor: JLA 16.250), for 30 min at 4°C. The cell pellets were then resuspended with 300 mL of buffer A (25 mM NaH_2_PO_4_, 25 mM Na_2_HPO_4_ pH 7.5, 300 mM NaCl, 20 mM Imidazole and 1 mM PMSF), frozen in liquid nitrogen and stored at −80°C.

All subsequent purification steps were performed at 4°C. The cell pellets were thawed and sonicated for 15 rounds on ice for 30 sec followed by a one min rest interval in each round. The lysate was centrifuged at 20000 rpm for 45 min in centrifuge Beckman Coulter Avanti^®^ J-E with a, JLA 25.50 rotor). The supernatant was collected and loaded into a 5 mL HisTrap FF crude column pre-equilibrated with buffer A. The column was then washed with 50 CVs of buffer B (25 mM NaH_2_PO_4_, 25 mM Na_2_HPO_4_ pH 7.5, 300 mM NaCl and 20 mM Imidazole) on a ÄKTA FPLC. Protein bound to the column was then eluted isocratically with 20 CVs of buffer C (25 mM NaH_2_PO_4_, 25 mM Na_2_HPO_4_ pH 7.5, 300 mM NaCl and 250 mM Imidazole). The purity of the fractions was monitored by SDS-PAGE (Invitrogen by Thermo Fisher Scientific) and then combined and quantified with Bradford reagent (BioRad, Solna, Sweden). A quarter of the protein was transferred into a SnakeSkin^®^ Dialysis membrane tubing (10 kDa) in 2 L of buffer E (10 mM HEPES pH 7.6, 300 mM NaCl) overnight for buffer exchange. The protein was harvested and concentrated to 18 mg/mL using an Amicon Ultra Centrifugal Filters (10 kDa) for subsequent MST assays.

The remaining three quarters of the protein were dialysed in a SnakeSkin® Dialysis membrane tubing supplied with 3 mM β-mercaptoethanol (β-ME) and Ulp1 protease (1 mg Ulp1 for 30 mg protein) in 2 L of buffer D (25 mM NaH_2_PO_4_, 25 mM Na_2_HPO_4_ pH 7.6, 300 mM NaCl and 3 mM β-ME) overnight to cleave the His-SUMO fusion protein. After dialysis, nickel-affinity purification was applied. The cleaved protein was loaded into the second pre-equilibrated HisTrap FF crude column followed by washing with 3 CVs of buffer B. The cleaved nsp1 in the flow through was collected. The purity and the concentration were verified by SDS-PAGE and Bradford reagent. Afterwards, the protein was concentrated to 25 mg/mL using an Amicon Ultra Centrifugal Filters (10 kDa), aliquoted into 100 μL fractions, frozen in liquid nitrogen and stored at −80°C for subsequent crystallisation.

#### Crystallisation of nsp1

SARS-CoV-2 nsp1_10-126_ crystallisation trials were conducted with sitting drop vapour diffusion using commercial crystallisation screens and the Mosquito^®^ crystallisation nanodrop robot (SPT Labtech, Melbourne, UK) on 96-well 3-drop plates (SWISSCI AG, Zug, Switzerland). The thawed SARS-CoV-2 nsp1_10-126_ was diluted to 20 mg/mL with buffer E. Each drop was set up by mixing 200 nL of SARS-CoV-2 nsp1_10-126_ with 200 or 400 nL of reservoir solution from the commercial crystallisation screens. The crystallisation plates were incubated at 18°C. Large crystals appeared after incubation overnight in several crystallising conditions in three different shapes (supplementary table S1).

Cryoprotectant solution was prepared by adding 10%, 15% or 20% DMSO into the precrystallisation buffer/ buffer E (10 mM HEPES and 300 mM NaCl). 200 nL of cryoprotectant was mixed with around 200 nL of protein drops with large crystals formed therein to give final DMSO concentrations of 5%, 7.5% and 10%. As three different shapes of crystals were observed, after incubation for 5 min at room temperature, large crystals of each type were harvested, cryo-cooled in liquid nitrogen, and stored in unipucks, which were also kept in liquid nitrogen ready for diffraction experiments.

Based on the X-ray diffraction results of the crystals, condition Index 44 (0.1 M HEPES pH 7.5, 25% w/v PEG3350) was chosen for further SARS-CoV-2 nsp1_10-126_ crystallisation in large quantities. Frozen stocks of SARS-CoV-2 nsp1_10-126_ were thawed on ice and centrifuged in a Thermo Scientific Pico 17 Microcentrifuge, 24-Pl Rotor at 4°C, 20000 rpm for 10 min to remove aggregates before the determination of the protein concentration. Subsequently, the protein stock was diluted to 20 mg/mL with precrystallisation buffer. 400 mL of Index condition 44 (0.1 M HEPES pH 7.5 and 25% w/v PEG3350) was added into each reservoir well. Five protein drops were set on each cover slip by mixing 1 μL of protein solution with 1 μL of the reservoir. The 24-well Linbro plates were placed at 18°C in an incubator.

#### Fragment soaking of nsp1 crystals

For nsp1 fragment soaking we used the same vapour diffusion protocol as described above. Fragments were selected from the Maybridge Ro3 library at a concentration of 200 mM dissolved in DMSO. Fragments were diluted in Index 44 solution to 40 mM containing 20% DMSO. An equal amount of DMSO was diluted in Index 44 solution to obtain 20% DMSO. Then, 1.5 μL of each fragment solutions were added into around 1 μL of crystallisation drops, making the final concentration of fragments 24 mM and approximately 12% DMSO. 1.5 μL of 20% DMSO solution was mixed with about 1 μL of crystallisation drops as protein-only samples, which were used to construct the ground state model required for PanDDA analysis. Drops were incubated at room temperature for 3-4 h followed by freezing of crystals in liquid nitrogen and stored in unipucks at the same temperature for subsequent diffraction experiments.

#### X-ray diffraction data collection, structure determination and refinement

For high-resolution data collection of SARS-CoV-2 nsp1_10-126_, protein crystals were soaked in 10% MPD for 15 sec and cryo-cooled in liquid nitrogen for subsequent data collection at ID30B. 2300 images were collected with 0.02 s exposure time, an oscillation of 0.05 degrees, and a beam size of 50 μm. Data collection of fragment-soaked nsp1_10-126_ crystals was conducted at beamline MASSIF-1[30] at the ESRF. The high-resolution diffraction data was indexed processed and integrated using the XDS package[54] using a minimum 0.5 value for I/sigma and statistically significant CC(1/2) values to define the high resolution cutoff. The diffraction data for fragment-soaked protein samples were automatically processed by various data reduction packages (autoPROC[55], autoPROC_staraniso[56], XDSAPP[57], XIA2_DIALS[56], EDNA_proc[58]) to generate merged and integrated data sets. The data sets with the best statistics in terms of resolution, completeness and merging statistics were downloaded from ISPyB. For the native high resolution structure, molecular replacement (MR) was performed in Phenix with the published coordinate set of SARS-CoV-2 nsp1_10-126_ (PDB entry 7K7P) as a search model. The obtained structure was first automatically refined in Phenix followed by manually inspection and real space refinement in COOT, and followed by at least a second round of refinement in Phenix. Several iterations of refinement in Phenix and visual inspection in COOT were performed until the R_free_ value did not decrease any further. The model derived from refinement against the highest resolution data set was used as the model in the following MR procedure for SARS-CoV-2 nsp1_10-126_-fragment complex data sets.

To identify fragment hits binding to SARS-CoV-2 nsp1_10-126_, PanDDA, a multi-dataset crystallographic analysis program was employed. Each dataset contains a coordinate file with the suffix “pdb” and an electronic density map file with the suffix “mtz” for each protein sample along with the chemical structure files with the suffix “pdb” and “cif” for the fragment in which the SARS-CoV-2 nsp1_10-126_ crystal was soaked. The “pdb” and “mtz” files were generated after MR by the program Dimple in CCP4[59] with the new model, while the chemical structure files for fragments were generated by Elbow in Phenix. In addition, 40 high-resolution datasets of native SARS-CoV-2 nsp1_10-126_ from the protein soaked in 12% DMSO solution were used to construct a “ground sate” model of SARS-CoV-2 nsp1_10-126_ in PanDDA.

After automatic analysis and inspection, the identified hits were verified manually in COOT by fitting the chemical structures of the fragments into their corresponding electron density. The complexes were then refined in Phenix to verify the correctness of the fragment conformations in binding events.

#### Thermal shift assays for nsp1-fragment complexes using nanoDSF

The protein concentration used in this assay was determined by analysing the unfolding curves of SARS-CoV-2 nsp1_10-126_ at different concentrations in the presence of 2% DMSO. Fragments dissolved in DMSO (200 mM stocks) were diluted in buffer E supplied with 0.05% Pluronic® F-127 and mixed with nsp1_10-126._ The final concentration of fragments was 4 mM containing 2% DMSO, while the final protein concentration was 1.25 mg/mL. After centrifugation for 5 min at 6000 rpm, the samples were loaded into Tycho NT.6 capillaries (TY-C001, NanoTemper, München, Germany) with 1.25 mg/mL SARS-CoV-2 nsp1_10-126_ in the presence of 2% DMSO as control and measured by the Tycho NT.6 instrument (NanoTemper, München, Germany) in triplicate. The resulting T_i_ values are presented as means ± SD (n = 3) and the △T_i_ values for each sample were calculated with the following equation.

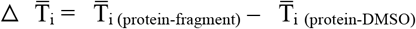

#### MST for the estimation of K_d_ values of fragment hits

To label the protein with fluorescent dye, 100 nM of RED-TRIS-NTA 2^nd^ Generation dye (MO-L018, NanoTemper, Müchen, Germany) was mixed with 800 nM SARS-CoV-2 nsp1_10-126_, SARS-CoV nsp1_2-180_ or MERS nsp1_2-193_ and incubated for 30 min on ice. The mixture was centrifuged in a Thermo Scientific Pico 17 MicroCentrifuge, 24-Pl Rotor at 15000 g for 10 min at 4°C to remove any aggregates. For fragment hits **5E11, 10B6**, and **11C6**, labelled protein was diluted to 200 nM in assay buffer (buffer E supplemented with 0.05% Pluronic(R) F-127) and the fragment hits were diluted from 200 mM to 20 mM with assay buffer. 20 μL of DMSO was mixed with 180 μL of assay buffer as ligand buffer. The fragment solution was serially diluted with ligand buffer with a dilution factor of two, obtaining 16 fragment solutions in gradient concentrations. Then, an equal volume of protein solution was mixed with each fragment solution and incubated for 30 min at 4°C to reach the binding equilibrium. The final fragment concentrations were from 10 mM to 305 nM with 5% DMSO. The final protein concentration in each sample was 100 nM. The samples were centrifuged at 6000 rpm for 5 min before the supernatant was being loaded into capillaries and detected in the Monolith NT.115 Pico instrument (NanoTemper, Müchen, Germany) under the Pico-RED channel with 5% excitation power and 40% MST power under the Binding Affinity mode in the MO. Control software. For fragment hits **7H2** and **8E6**, labelled protein was diluted to 80 nM in assay buffer and the fragment hits were serially diluted from 200 mM to 40 mM with assay buffer. 40 μL of DMSO was mixed with 160 μL of assay buffer as ligand buffer. The fragment solution was serially diluted with ligand buffer with a dilution factor of 1.3, obtaining 16 fragment solutions in gradient concentrations. Then, an equal volume of protein solution was mixed with each fragment solution and incubated for 30 min at 4°C to reach the binding equilibrium. The final fragment concentrations were from 20 mM to 390 nM with 5% DMSO. The final protein concentration in each sample was 40 nM. The samples were centrifuged at 6000 rpm for 5 min before the supernatant was loaded into capillaries and detected under the Pico-RED channel with 20% excitation power and 40% MST power under the Binding Affinity mode in the MO. Control software. The temperature was set at 25°C. Each fragment hit was tested at least in triplicate. The data were analysed, and the figures were generated in the MO. Affinity Analysis software.

#### Calculation of physico-chemical properties and preparation of figures

The molecular weight (MW), the clogP and the polar surface area (PSA) of fragment hits were calculated from the chemical structures using ChemDraw version 19.1. Hydrogen-bond donors and acceptors as well as clogS were calculated on the drug-likeness prediction server of Molsoft: http://molsoft.com/mprop/. Figures were prepared using ChemDraw 19.2, Pymol (The PyMOL Molecular Graphics System, Version 2.4.1, Schrödinger, LLC), Chimera[60] and Liplot+[61]. The default values for hydrogen-bond interactions (2.7 to 3.35 Å) and non-bonded contacts (2.9 to 3.9 Å) were used.

## Supporting information

Supplementary material section

## Abbreviations

CoV: Coronavirus
EDTA: Ethylenediaminetetraacetic Acid;
IPTG: Isopropyl-d-thiogalactopyranoside
MERS: Middle East Respiratory Syndrome
NSP: Nonstructural protein
ORF: Open Reading Frame
PMSF: Phenylmethylsulfonylfluorid
SDS-PAGE: Sodium Dodecyl Sulphate–polyacrylamide Gel Electrophoresis
SARS: Severe Acute Respiratory Syndrome
CoV: Coronaviruses
SARS-CoV: Severe Acute Respiratory Syndrome Coronavirus
MERS-CoV: Middle East Respiratory Syndrome Coronavirus;
SAR: Structure activity relationship
IFN: Interferon
MR: Molecular replacement
GS: Ground state
DMSO: Dimethyl Sulfoxide
HEPES: 4-(2-hydroxyethyl)-1-piperazineethanesulfonic Acid
LB: Terrific Broth
TB: Terrific Broth
SOC: Super Optimal Broth
MST: Microscale Thermophoresis
TSA: Thermal Shift Assay
SUMO: Small Ubiquitin-like Modifier
His-tag: Polyhistidine-tag
β-ME: β-mercaptoethanol
ULP1: Ubiquitin-like-specific Protease 1
IPTG: Isopropylthio-β-galactoside
PDB: Protein Data Bank
COOT: Crystallographic Object-oriented Toolkit
PanDDA: Pan-Dataset Density Analysis
NanoDSF: Nanoscale Differential Scanning Fluorimetry
PEG: Polyethylene Glycol
PBS-T: 1X Phosphate-buffered Saline with 0.1% Tween
*E. coli*: *Escherichia coli*
SD: Standard Deviation
SL: Stem Loop
Pol α: DNA Polymerase α.

## Author contributions

SM, SD, JL, NP, MB, SH, FK

## Acknowledgements

Fragment-based screening for nsp1 via X-ray crystallography was conducted at MASSIF-1 beamline at the ESRF, Grenoble, France. We thank members of at the ESRF, beamline ID30B and at DLS, Oxfordshire, UK, beamline I04, for support collecting atomic-resolution data for SARS-CoV-2 nsp1. This manuscript contains part of Shumeng Ma’s PhD work.

## References

1. Zhong, N.S., et al., Epidemiology and cause of severe acute respiratory syndrome (SARS) in Guangdong, People’s Republic of China, in February, 2003. The Lancet, 2003. 362(9393): p. 1353–1358.

2. Zaki, A.M., et al., Isolation of a novel coronavirus from a man with pneumonia in Saudi Arabia. N Engl J Med, 2012. 367(19): p. 1814–20.

3. Xiao, X., et al., Animal sales from Wuhan wet markets immediately prior to the COVID-19 pandemic. Sci Rep, 2021. 11(1): p. 11898.

4. Zhao, Y.J., et al., Mental health status and quality of life in close contacts of COVID-19 patients in the post-COVID-19 era: a comparative study. Transl Psychiatry, 2021. 11(1): p. 505.

5. Sen, P., et al., Burden and characteristics of COVID-19 in the United States during 2020. Nature, 2021.

6. Josephson, A., T. Kilic, and J.D. Michler, Socioeconomic impacts of COVID-19 in low-income countries. Nat Hum Behav, 2021. 5(5): p. 557–565.

7. Salje, H., et al., Estimating the burden of SARS-CoV-2 in France. Science, 2020. 369(6500): p. 208–211.

8. Aruffo, E., et al., Community structured model for vaccine strategies to control COVID19 spread: a mathematical study. medRxiv, 2021: p. 2021.01.25.21250505.

9. Porter, J.R., Vaccines, social measures and Covid19 - A European evidence-based analysis Vaccines, social measures and Covid19. medRxiv, 2021: p. 2021.04.15.21255558.

10. Lipsitch, M. and N.E. Dean, Understanding COVID-19 vaccine efficacy. Science, 2020. 370(6518): p. 763–765.

11. Madison, A.A., et al., Psychological and behavioral predictors of vaccine efficacy: Considerations for COVID-19. Perspectives on Psychological Science, 2021. 16(2): p. 191–203.

12. Cai, Y., et al., Structural basis for enhanced infectivity and immune evasion of SARS-CoV-2 variants. bioRxiv, 2021.

13. Abu-Raddad, L.J., H. Chemaitelly, and A.A. Butt, Effectiveness of the BNT162b2 Covid-19 Vaccine against the B. 1.1. 7 and B. 1.351 Variants. New England Journal of Medicine, 2021.

14. The Pfizer BioNTech (BNT162b2) COVID-19 vaccine: What you need to know. 2 September 2021; Available from: https://www.who.int/news-room/feature-stories/detail/who-can-take-the-pfizer-biontech-covid-19--vaccine?adgroupsurvey={adgroupsurvey}&gclid=EAIaIQobChMIqpiuz4u08wIVkOmyCh0UlQ2KEAAYASABEgIIxvD_BwE.

15. Parvathaneni, V. and V. Gupta, Utilizing drug repurposing against COVID-19 - Efficacy, limitations, and challenges. Life Sci, 2020. 259: p. 118275.

16. Zhou, P., et al., A pneumonia outbreak associated with a new coronavirus of probable bat origin. Nature, 2020. 579(7798): p. 270–273.

17. Masters, P.S., The molecular biology of coronaviruses. Advances in virus research, 2006. 66: p. 193–292.

18. Min, Y.Q., et al., SARS-CoV-2 nsp1: Bioinformatics, Potential Structural and Functional Features, and Implications for Drug/Vaccine Designs. Front Microbiol, 2020. 11: p. 587317.

19. Narayanan, K., et al., Severe acute respiratory syndrome coronavirus nsp1 suppresses host gene expression, including that of type I interferon, in infected cells. J Virol, 2008. 82(9): p. 4471–9.

20. Terada, Y., et al., MERS coronavirus nsp1 participates in an efficient propagation through a specific interaction with viral RNA. Virology, 2017. 511: p. 95–105.

21. Lapointe, C.P., et al., Dynamic competition between SARS-CoV-2 NSP1 and mRNA on the human ribosome inhibits translation initiation. Proc Natl Acad Sci U S A, 2021. 118(6).

22. Thoms, M., et al., Structural basis for translational shutdown and immune evasion by the Nsp1 protein of SARS-CoV-2. Science, 2020. 369(6508): p. 1249–1255.

23. Schubert, K., et al., SARS-CoV-2 Nsp1 binds the ribosomal mRNA channel to inhibit translation. Nat Struct Mol Biol, 2020. 27(10): p. 959–966.

24. Shi, M., et al., SARS-CoV-2 Nsp1 suppresses host but not viral translation through a bipartite mechanism. bioRxiv, 2020.

25. Zhang, K., et al., Nsp1 protein of SARS-CoV-2 disrupts the mRNA export machinery to inhibit host gene expression. Science Advances, 2021. 7(6): p. eabe7386.

26. Lokugamage, K.G., et al., Middle East Respiratory Syndrome Coronavirus nsp1 Inhibits Host Gene Expression by Selectively Targeting mRNAs Transcribed in the Nucleus while Sparing mRNAs of Cytoplasmic Origin. J Virol, 2015. 89(21): p. 10970–81.

27. Semper, C., N. Watanabe, and A. Savchenko, Structural characterization of nonstructural protein 1 from SARS-CoV-2. iScience, 2021. 24(1): p. 101903.

28. Alonso, G., et al., Differential Activation of p38 Mitogen-activated Protein Kinase Isoforms Depending on Signal Strength*. Journal of Biological Chemistry, 2000. 275(51): p. 40641–40648.

29. Svensson, O., et al., Multi-position data collection and dynamic beam sizing: recent improvements to the automatic data-collection algorithms on MASSIF-1. Acta Crystallographica Section D, 2018. 74(5): p. 433–440.

30. Bowler, M.W., et al., MASSIF-1: a beamline dedicated to the fully automatic characterization and data collection from crystals of biological macromolecules. Journal of Synchrotron Radiation, 2015. 22(6): p. 1540–1547.

31. Pearce, N.M., et al., A multi-crystal method for extracting obscured crystallographic states from conventionally uninterpretable electron density. Nature Communications, 2017. 8(1): p. 15123.

32. Emsley, P., et al., Features and development of Coot. Acta Crystallographica Section D: Biological Crystallography, 2010. 66(4): p. 486–501.

33. Afonine, P.V., et al., Towards automated crystallographic structure refinement with phenix.refine. Acta Crystallographica Section D, 2012. 68(4): p. 352–367.

34. Kozielski, F., et al., Identification of fragments binding to SARS-CoV-2 nsp10 reveals ligand-binding sites in conserved interfaces between nsp10 and nsp14/nsp16. RSC Chemical Biology, 2022. 3(1): p. 44–55.

35. Abagyan, R., M. Totrov, and D. Kuznetsov, ICM—A new method for protein modeling and design: Applications to docking and structure prediction from the distorted native conformation. Journal of Computational Chemistry, 1994. 15(5): p. 488–506.

36. Thompson, J.D., D.G. Higgins, and T.J. Gibson, CLUSTAL W: improving the sensitivity of progressive multiple sequence alignment through sequence weighting, position-specific gap penalties and weight matrix choice. Nucleic acids research, 1994. 22(22): p. 4673–4680.

37. Robert, X. and P. Gouet, Deciphering key features in protein structures with the new ENDscript server. Nucleic Acids Research, 2014. 42(W1): p. W320–W324.

38. Vankadari, N., N.N. Jeyasankar, and W.J. Lopes, Structure of the SARS-CoV-2 Nsp1/51-Untranslated Region Complex and Implications for Potential Therapeutic Targets, a Vaccine, and Virulence. The Journal of Physical Chemistry Letters, 2020. 11(22): p. 9659–9668.

39. Gordon, D.E., et al., A SARS-CoV-2 protein interaction map reveals targets for drug repurposing. Nature, 2020. 583(7816): p. 459–468.

40. Pellegrini, L., The Pol α-Primase Complex, in The Eukaryotic Replisome: a Guide to Protein Structure and Function, S. MacNeill, Editor. 2012, Springer Netherlands: Dordrecht. p. 157–169.

41. Starokadomskyy, P., A. Escala Perez-Reyes, and E. Burstein, Immune Dysfunction in Mendelian Disorders of POLA1 Deficiency. Journal of Clinical Immunology, 2021. 41(2): p. 285–293.

42. Starokadomskyy, P., et al., DNA polymerase-α regulates the activation of type I interferons through cytosolic RNA:DNA synthesis. Nature Immunology, 2016. 17(5): p. 495–504.

43. Kilkenny, M.L., et al., Structural basis for the interaction of SARS-CoV-2 virulence factor nsp1 with DNA polymerase alpha-primase. Protein Sci, 2022. 31(2): p. 333–344.

44. Mou, K., et al., Emerging Mutations in Nsp1 of SARS-CoV-2 and Their Effect on the Structural Stability. Pathogens (Basel), 2021. 10(10): p. 1285.

45. Enslen, H., D.M. Brancho, and R.J. Davis, Molecular determinants that mediate selective activation of p38 MAP kinase isoforms. The EMBO Journal, 2000. 19(6): p. 1301–1311.

46. Kumar, A., et al., SARS-CoV-2 NSP1 C-terminal (residues 131–180) is an intrinsically disordered region in isolation. Current Research in Virological Science, 2021. 2: p. 100007.

47. Schubert, K., et al., SARS-CoV-2 Nsp1 binds the ribosomal mRNA channel to inhibit translation. Nature Structural & Molecular Biology, 2020. 27(10): p. 959–966.

48. Mendez, A.S., et al., The N-terminal domain of SARS-CoV-2 nsp1 plays key roles in suppression of cellular gene expression and preservation of viral gene expression. Cell Rep, 2021. 37(3): p. 109841.

49. Kamitani, W., et al., Severe acute respiratory syndrome coronavirus nsp1 protein suppresses host gene expression by promoting host mRNA degradation. Proceedings of the National Academy of Sciences, 2006. 103: p. 12885–12890.

50. Kamitani, W., et al., A two-pronged strategy to suppress host protein synthesis by SARS coronavirus Nsp1 protein. Nature Structural & Molecular Biology, 2009. 16(11): p. 1134–1140.

51. Huang, C., et al., SARS Coronavirus nsp1 Protein Induces Template-Dependent Endonucleolytic Cleavage of mRNAs: Viral mRNAs Are Resistant to nsp1-Induced RNA Cleavage. PLOS Pathogens, 2011. 7(12): p. e1002433.

52. Shi, M., et al., SARS-CoV-2 Nsp1 suppresses host but not viral translation through a bipartite mechanism. bioRxiv : the preprint server for biology, 2020: p. 2020.09.18.302901.

53. Shuvalov, A., et al., Nsp1 of SARS-CoV-2 stimulates host translation termination. RNA Biology, 2021. 18(sup2): p. 804–817.

54. Kabsch, W., XDS. Acta Crystallographica Section D, 2010. 66(2): p. 125–132.

55. Vonrhein, C., et al., Data processing and analysis with the autoPROC toolbox. Acta Crystallographica Section D, 2011. 67(4): p. 293–302.

56. Vonrhein, C., et al., Advances in automated data analysis and processing within autoPROC, combined with improved characterisation, mitigation and visualisation of the anisotropy of diffraction limits using STARANISO. Acta Crystallographica Section A, 2018. 74(a1): p. a360.

57. Krug, M., et al., XDSAPP: a graphical user interface for the convenient processing of diffraction data using XDS. Journal of Applied Crystallography, 2012. 45(3): p. 568–572.

58. Incardona, M.-F., et al., EDNA: a framework for plugin-based applications applied to X-ray experiment online data analysis. Journal of synchrotron radiation, 2009. 16(6): p. 872–879.

59. Winn, M.D., et al., Overview of the CCP4 suite and current developments. Acta crystallographica. Section D, Biological crystallography., 2011. 67(4): p. 235–242.

60. Pettersen, E.F., et al., UCSF Chimera--a visualization system for exploratory research and analysis. Journal of computational chemistry, 2004. 25(13): p. 1605–12.

61. Laskowski, R.A. and M.B. Swindells, LigPlot+: Multiple Ligand–Protein Interaction Diagrams for Drug Discovery. Journal of Chemical Information and Modeling, 2011. 51(10): p. 2778–2786.

