## Supplementary material section for "Two ligand-binding sites on SARS-CoV-2 non-structural protein 1 revealed by fragment-based x-ray screening"

### Content

Table S1. Crystal images with crystallisation conditions 2

Table S2. Data collection and refinement statistics for SARS-CoV-2 $nsp1_{10\text{-}126}$-fragment complexes 3

**Table S1:** Representative crystallisation conditions and images of three types of SARS-CoV-2 nsp1$_{10\text{-}126}$ crystals. All crystals grew at 18°C at protein concentrations between 15 and 27 mg/ml.

| **Commercial screens** | **Crystallisation condition** | **Crystal images** |
|---|---|---|
| Index™ | 0.1 HEPES pH 7.5, 25% w/v PEG3350 | 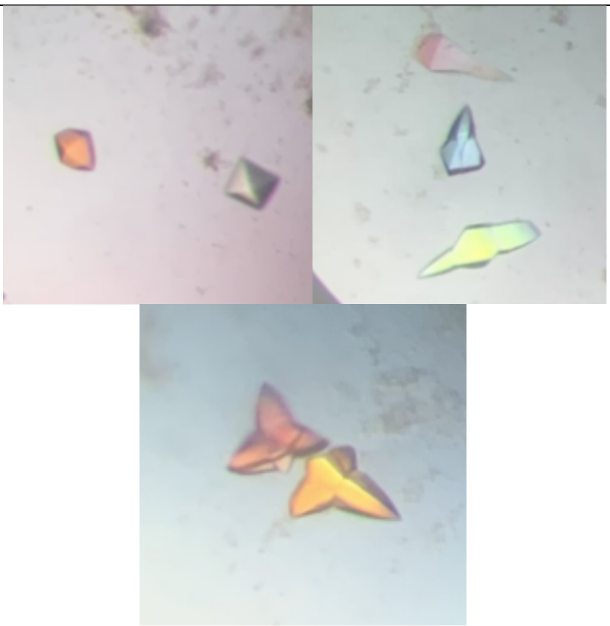 |
| Morpheus® | 0.12 M Ethylene glycol, 0.1 M Imidazole, 0.1 M MES monohydrate acid pH 6.5, 20% v/v Ethylene glycol, 10% w/v PEG8000 | |
| JCSG-*plus*™ | 0.2 M Ammonium formate, 20% w/v PEG3350 | |

**Table S2.** Data collection, data processing, and model refinement statistics for four nsp1$_{10-126}$-fragment complexes from SARS-CoV-2. Data in parenthesis correspond to the highest resolution shell.

| | **Nsp1-11C6** | **Nsp1-10B6** | **Nsp1-5E11** |
|---|---|---|---|
| **PDB ID** | xxx | xxx | xxx |
| **Wavelength [Å]** | 0.9655 | 0.9655 | 0.9655 |
| **Resolution range [Å]** | 35.63-1.28 (1.36-1.28) | 46.99-1.37 (1.39 - 1.37) | 47.16-1.04 (1.10-1.04) |
| **Space group** | $P4_32_12$ | $P4_32_12$ | $P4_32_12$ |
| **Unit cell parameters (Å, °)** | *a*=*b*=36.817, *c*=141.205, α=β=γ=90 | *a*=*b*=36.697, *c*=140.96, α=β=γ=90 | *a*=*b*=36.705, *c*=141.491, α=β=γ=90 |
| **Total reflections** | 162517 (9876) | 192744 (1644) | 357252 (19766) |
| **Unique reflections** | 22326 (1116) | 19918 (619) | 83829 (10752) |
| **Multiplicity** | 7.28(8.85) | 9.68 (2.66) | 4.26 (1.84) |
| **Completeness [%]** | 92.9 (46.9) | 93.7 (60.5) | 94.3 (74.8) |
| **Mean I/sigma(I)** | 15.8 (1.5) | 23.7 (1.1) | 5.69 (0.12) |
| **Wilson B-factor ($Å^2$)** | 19.6 | 14.2 | 11.8 |
| **R-meas** | 0.06 (1.26) | 0.053 (0.221) | 0.10 (5.56) |
| **$CC_{1/2}$** | 1.0 (0.65) | 1.0 (0.92) | 1.0 (0.52) |
| **$R_{cryst}/R_{free}$ [%]** | 14.9 (21.1) / 20.6 (28.0) | 15.2 (15.9) / 20.9 (25.8) | 21.3 (65.6) / 25.4 (67.7) |
| **Total no. of non-hydrogen atoms (protein)** | 1152 | 1140 | 994 |
| **No. of protein/ligand/solvent atoms** | 1013 / 15 / 124 | 1006 / 26 / 134 | 897 / 14 / 83 |
| **Average B-factor/protein/ligands/solvent** | 24.4 / 23/1 / 46.7 / 32.6 | 20.8 / 18.9 / 31.3 / 34.7 | 25.3 / 23.7 / 72.1 / 34.6 |
| **RMSD (bonds, angles)** | 0.012 / 1.34 | 0.012 / 1.37 | 0.012 / 1.44 |
| **Ramachandran favored/allowed/outliers/rotamer outliers [%]** | 97.3 / 2.7 / 0.0 / 0.9 | 97.3 / 2.7 / 0.0 / 0.0 | 98.3 / 1.7 / 0.0 / 1.0 |
| **Clashscore** | 4.4 | 4.6 | 3.8 |

**Continuation Table S2.** Data collection, data processing, and model refinement statistics for four nsp1$_{10\text{-}126}$-fragment complexes from SARS-CoV-2. Data in parenthesis correspond to the highest resolution shell.

| | **Nsp1-8E6** | **Nsp1-7H2** |
|---|---|---|
| **PDB ID** | xxx | xxx |
| **Wavelength [Å]** | 0.9655 | 0.9655 |
| **Resolution range [Å]** | 35.47 - 1.20 (1.18-1.18) | 36.52-1.04 (1.10-1.04) |
| **Space group** | $P4_32_12$ | $P4_32_12$ |
| **Unit cell parameters (Å, °)** | *a*=*b*=36.643, *c*=141.366, α=β=γ=90 | *a*=*b*=36.524, *c*=141.988, α=β=γ=90 |
| **Total reflections** | 297854 (15197) | 483869 (27003) |
| **Unique reflections** | 30526 (1578) | 85622 (11714) |
| **Multiplicity** | 9.8 (9.6) | 5.65 (2.31) |
| **Completeness [%]** | 92.8 (100) | 96.9 (81.7) |
| **Mean I/sigma(I)** | 18.3 (2.2) | 10.18 (0.40) |
| **Wilson B-factor (Å$^2$)** | 15.3 | 12.2 |
| **R-meas** | 0.06 (1.02) | 0.075 (2.125) |
| **$CC_{1/2}$** | 1.00 (0.80) | 1.00 (0.17) |
| **$R_{cryst}/R_{free}$ [%]** | 17.1 (19.8) / 21.2 (25.7) | 17.6 (45.1) / 20.6 (47.5) |
| **Total no. of non-hydrogen atoms (protein)** | 970 | 1032 |
| **No. of protein/ligand/solvent atoms** | 878 / 11 / 81 | 919 / 10 / 103 |
| **Average B-factor/protein/ligands/solvent** | 21.1 / 20.5 / 27.7 / 26.3 | 23.7 / 22.3 / 57.4 / 33.3 |
| **RMSD (bonds, angles)** | 0.013 / 1.52 | 0.012 / 1.41 |
| **Ramachandran favored/allowed/outliers/rotamer outliers [%]** | 97.4 / 2.6 / 0.0 / 0.0 | 95.6 / 3.5 / 0.9 / 1.0 |
| **Clashscore** | 2.3 | 6.9 |
